# Dose-dependent interaction of parasites with tiers of host defence generates “wormholes” that prolong infection at intermediate inoculum sizes

**DOI:** 10.1101/2023.06.09.544354

**Authors:** Andrea L. Graham, Roland R. Regoes

## Abstract

Immune responses are induced by parasite exposure and can in turn reduce parasite burden. Despite such apparently simple rules of engagement, key drivers of within-host dynamics, including dose-dependence of defence and infection duration, have proven difficult to predict. Here, we model how varied inoculating doses interact with host defences. Defence is multi-tiered, with 3 tiers invoked for all animals: barrier, innate, and adaptive. We model how these tiers interact with replicating and non-replicating parasites across six orders of magnitude of dose. We find that, in general, intermediate parasite doses generate infections of longest duration because they are sufficient in number to breach barrier defences, but insufficient to strongly induce subsequent tiers of defence. Deviation from the hypothesis of independent action, which postulates that each parasite has an independent probability of establishing infection, may therefore be widespread. Most interestingly, our model predicts local maxima of duration at two doses – one for each tier transition. While empirical evidence is consistent with nonlinear dose-dependencies, the profiles with multiple turning points that we predict will require finer-scale dose experiments than are usually undertaken. Our results help explain varied infection duration among differentially-exposed hosts and elucidate evolutionary pressures that shape both virulence and defence.

## Introduction

Following exposure to parasites, the dynamics of parasite growth and immune response induction are important determinants of infectivity (probability of establishing infection) as well as severity and chronicity of infection. For example, hosts that exhibit extremely rapid induction and decay of immune responses are expected to clear infection quickly while minimizing costs of defence (1). However, the dose of parasites to which a host is exposed is likely to alter these within-host dynamics, both qualitatively and quantitatively: inoculating dose may determine whether infection is established at all (2-6) as well as the duration or severity of infections that do establish (7-10). General principles governing dose-dependence of infection and immunity therefore merit formal investigation.

A prevalent and useful conceptualisation of the quantitative process of infection is the hypothesis of independent action. According to this hypothesis, each individual parasite (e.g., virus, bacterium, fungus, protozoan, helminth, or ectoparasite) has the same small chance to initiate an infection, and this chance is independent of how many other parasites are present in the inoculum (11, 12). Because each parasite can initiate an infection independently if the hypothesis of independent action holds, the cumulative infection probability is predicted to increase monotonically with the inoculum dose (12). Many systems have been found to conform to the hypothesis of independent action, including classic work on *Haemophilus influenzae* in rats (13) (but see (14)), as well as diverse virus infections in plants (4), insects (3), and fish (5)).

Deviations from the independent action hypothesis have also been conceived and documented. Halvorson (11), who first formulated the hypothesis of independent action, contrasted it with a hypothesis that a critical number of parasites is required for infection. Requiring a critical, or threshold, number is not consistent with parasite individuals acting independently. Rather it constitutes a form of synergy – e.g., when a quorum of bacteria in insects (15) or a quorum of bacteria (16) or protozoa (17) in mammals must cooperatively signal or differentiate to achieve or sustain infection. Mechanistically, such cooperativity could arise directly by signaling among individual parasites, or indirectly (14), for example as the result of the interaction between the parasite population and the immune system of the host (focal to this study; see below).

Another apparent deviation from independent action arises when hosts differ in susceptibility (e.g., due to variation among individuals in rates of immune response induction). In this case, the increase in infectivity with inoculum dose is flatter than predicted by the hypothesis of independent action, and this slope can be used to quantify the variation in susceptibility among hosts in the population (18-23). This apparent deviation does not require a lack of independence of parasites during infection, but arises from the population-level effects of susceptibility differences. Such explanations for the shape of the relationship between inoculum dose and infection success, though informative, do not fully account for the essence of the host-parasite interaction: e.g., the reciprocal quantitative dependence of immune response induction on parasite abundance and of parasite abundance on immune responses. Indeed, even though non-independent action has been invoked as a contrast to independent action, the interactive, mechanistic basis of dose-dependence has rarely been clarified (24). A notable exception is the study by Pujol et al. (2), in which cooperativity arose as a consequence of the interaction between the parasite population and a single-tiered immune response. Pujol et al. developed a model in which the parasite elicited an immune effector that curbed its growth. Allowing such feedbacks generated complex relationships between inoculum dose and infection success that were consistent with data on poliovirus, among other infections (2). The interval of time over which parasites were inoculated was crucial to their results, which is perhaps unsurprising, given the time-dependence of immune processes as well as parasite replication.

Here, we formulate mathematical models to investigate how inoculating dose interacts with multi-tiered immune defences to determine duration of infection, including whether or not infection establishes at all. Like Pujol (2) and a few others (25, 26), we thus go beyond the simplification of the classic independent action hypothesis, and mathematically capture more realistic interplay between the parasites and the immune system within the host. We consider that not only do parasites induce immune responses (roughly according to “mass action” of the rates of encounter between parasites and immune cells (27)), but also that different immune system components differ in how and when they are triggered, when they are expressed, and how effective they are (following, e.g., (28, 29)).

Specifically, we model host defences that are multi-tiered, with barrier, innate, and adaptive defences as described in mammals (e.g., (30, 31)), among other animals. Physical and chemical barrier defences are constitutive (like skin) or maintained at steady-state levels (like antimicrobial peptides or mucosal antibodies). Such defences are immediately ready upon exposure but can, in principle, be overcome or eroded at sufficiently high inoculating doses. Next, non-specific innate defences like macrophages consuming microbes are rapidly induced (within hours) and subject to handling time saturation (such that large numbers of parasites can overwhelm them (32, 33)). Finally, specific adaptive defences like killer T cell or antibody responses are induced more slowly (days to weeks (31)) but are capable of achieving extremely high concentrations that might conquer large numbers of parasites.

Our model based on these complexities predicts relationships for the establishment and chronicity of infection with inoculum dose that are inconsistent with the hypothesis of independent action. The inconsistencies we report are more dramatic than in previous studies: not only do we predict that infection success and duration do not increase linearly with dose, but we find profiles with multiple peaks at intermediate parasite doses. The fact that infection success may not simply increase monotonically with inoculum dose has important implications for the estimation of host susceptibility, for epidemiology, and for the evolution of strategies for both attack and defence. The prediction of multiple peaks in infection success and duration, if empirically confirmed with future dose-response studies, could be exploited to determine the number of immune response tiers that are functional in a specific host-parasite system.

### Model definition

We developed a model that describes the growth of the parasite population within the host, and its inhibition by the immune system. We assume no anatomical structure within the host; the population dynamics therefore unfolds in a single, well-mixed compartment.

Let, *P* denote the number of parasites in this compartment. Parasite population dynamics are described by the following differential equation:

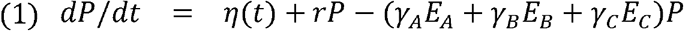

Hereby *η*(*t*) is a time-dependent function that captures the exposure of the host. The integral over this function, 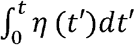, is the total number of parasites that enter the host, either through natural exposure or experimental inoculation. Because, in this study, we are focusing on the implications of dosing on the within host dynamics, this function is central.

The form of the function *η*(*t*) allows very flexible dosing schedules. Inoculations with a single dose (“bolus”) can be described by function with a high short peak for a short duration, where the height multiplied by the duration gives the overall dose. Alternatively, continuous (“trickle”) or repeated inoculations can also be described.

Once the host has been seeded with parasites, they start to replicate exponentially at a rate *rP* and are inhibited by three immune effectors, the number of which we denote by *E*_*A*_, *E*_*B*_, and *E*_*C*_. These three immune effector tiers, rather than describing specific actors that can differ among infections or host types, cover the relevant dynamical range of immune responses. In particular, we consider a constitutively expressed effector *E*_*A*_, and two inducible effectors, *E*_*B*_ and *E*_*C*_, that differ in the speed at which they are induced.

The slower one, *E*_*C*_, is ultimately more effective.

The dynamics of these immune effectors is governed by the following equations:

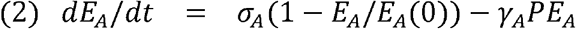

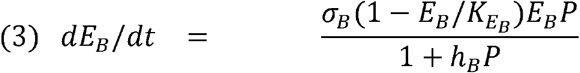

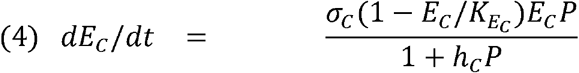

*E*_*A*_ is constitutively expressed at the level *E*_*A*_(0). This effector is depleted when it kills the parasite at a rate *y*_*A*_*PE*_*A*_. It is replenished at a maximum rate *a*_*A*_. The other two effectors. *E*_*B*_ and *E*_*C*_, are induced at rates that depend on the parasite load *P*. At low parasite loads, *P* and low levels of *E*_*B*_ and *E*_*C*_, they are induced at exponential rates *σ*_*B*_*P* and *σ*_*C*_*P*, respectively. The parameters *h*_*B*_ or *h*_*C*_ denote the parasite loads at which these two rates are half the maximum. The induction rate goes to zero if the levels of the immune effectors reach their respective carrying capacities 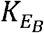 and 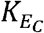. Unlike *E*_*A*_, *E*_*B*_ and *E*_*C*_ are not assumed to be decimated by killing parasites. The dynamics of *E*_*B*_ and *E*_*C*_ are structurally identical, but they differ in terms of the speed of induction (a_B_ and a_C_) and maximum efficacy (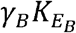 and 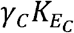)

### Model parameterisation and scenarios

The model is parameterized generically. This means that, rather than simulating a specific infection, we choose our parameters such that they recapitulate the typical time scales of the generation of the three tiers of immune responses upon parasite exposure.

In the first instance, we assumed that, following arrival at an array of inoculating doses, the parasite replicates at a per capita rate of 1 per day, which corresponds to a doubling time of approximately 0.7 days, or a 10-fold increase in approximately 2.3 days. Parasite growth is not assumed to be limited by a carrying capacity. When we instead simulate macroparasitic infection, we assume that parasites enter the hosts at various numbers, but do not replicate. We also ran simulations for singular (“bolus”) or continuous (“trickle”) exposure to both micro- and macroparasites.

The first tier of the immune system is constitutively expressed at the arbitrarily chosen level of 100 effectors, *E*_*A*_. As for mucosal antibodies, by exerting their effect, they are depleted at a rate that depends on the inoculum size, leading to either an immediate depletion of the effectors when facing a large inoculum dose, or to a fast clearance of a small inoculum. Once the parasites are cleared, these first-tier effectors grow back to the initial level of 100 at an initial rate of *σ*_*A*_ = 30 per day. The rate is reduced linearly until the homeostatic level is reached.

The second tier of the immune system, *E*_*B*_, is conceived as innate immune components that are initially set to a single effector. They are stimulated by parasites that overcome the first-tier response to proliferate/recruit at a logistic rate that depends on the parasite load. A single second-tier effector is assumed to be slightly less effective than a single first-tier effector. These second tier effectors can reach a carrying capacity of 10’000. At this level they are still assumed to be 20% less potent than the barrier response at its constitutively expressed level, to capture handling time constraints for effectors such as macrophages (32, 33) compared to mucosal antibodies at the barrier (34, 35). Unlike the first-tier effectors, however, second-tier effectors are not depleted by exerting their effect. Thus, the second-tier is effective across a larger range of parasite doses.

The third-tier of the immune system, *E*_*C*_, is conceived as adaptive immune effectors that are assumed to differ from the second tier only quantitatively, in terms of both a slower induction rate and higher effector efficacy. They are elicited to proliferate by the parasites that overcome the first-tier responses at a rate that is two orders of magnitude lower than the induction rate of the second tier. Their per-effector potency to clear parasites is assumed to be the same as that of the first-tier effectors. Their carrying capacity is set to 10^6^ but in practice they stop proliferating before that cap because they typically lead to fast parasite clearance.

We deviate from this default parameterisation to investigate the infection dynamics in immune-tier “knockout” hosts. These tier “knockouts” are implemented by setting the initial concentration of the respective immune effectors, *E*_*A*_, *E*_*B*_, or *E*_*C*_, to zero. This procedure mirrors knocking out genes responsible for various immune components in mice in experimental immunology (36), but is, due to our conceptual approach, more generic and cleaner because it is without pleiotropic effects on other traits.

### Model implementation

We implemented the deterministic population dynamical model in Equations 1-4 in the R language for statistical computing (37) using the function lsoda in the package deSolve (38).

We also implemented a stochastic version of the model using the implementation of the Gillespie algorithm in the R-package adaptivetau (39) that implements the adaptive tau-leaping approximation for simulating the trajectory of a continuous-time Markov (40). For the stochastic implementation, the logistic growth terms in Equations 3 and 4 have been partitioned into two terms, on describing the population expansion (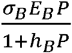and 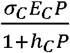), the other describing death due to crowding (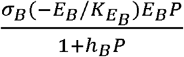and 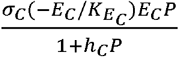). (This partitioning of the growth terms is generally required for the stochastic implementation of logistic growth to prevent the population turnover to be zero at the carrying capacity.)

For all details on our simulation models and the parameterisation please refer to code that is provided as supplementary online material.

## Results

We studied the dose-dependence of immune response induction and within-host dynamics of infection using a mathematical model. The model describes the how the parasite invades the host, replicates in it, and is confronted with three tiers of defence – roughly corresponding to the constitutively maintained barrier, the induced innate, and the induced adaptive tiers. Initially, we ran deterministic model simulations varying the inoculum size (dose) across fourorders of magnitude. Parasites and the three immune components exhibited diverse dose-dependent dynamics (**Figure 1**), as follows.

**Figure 1A:**
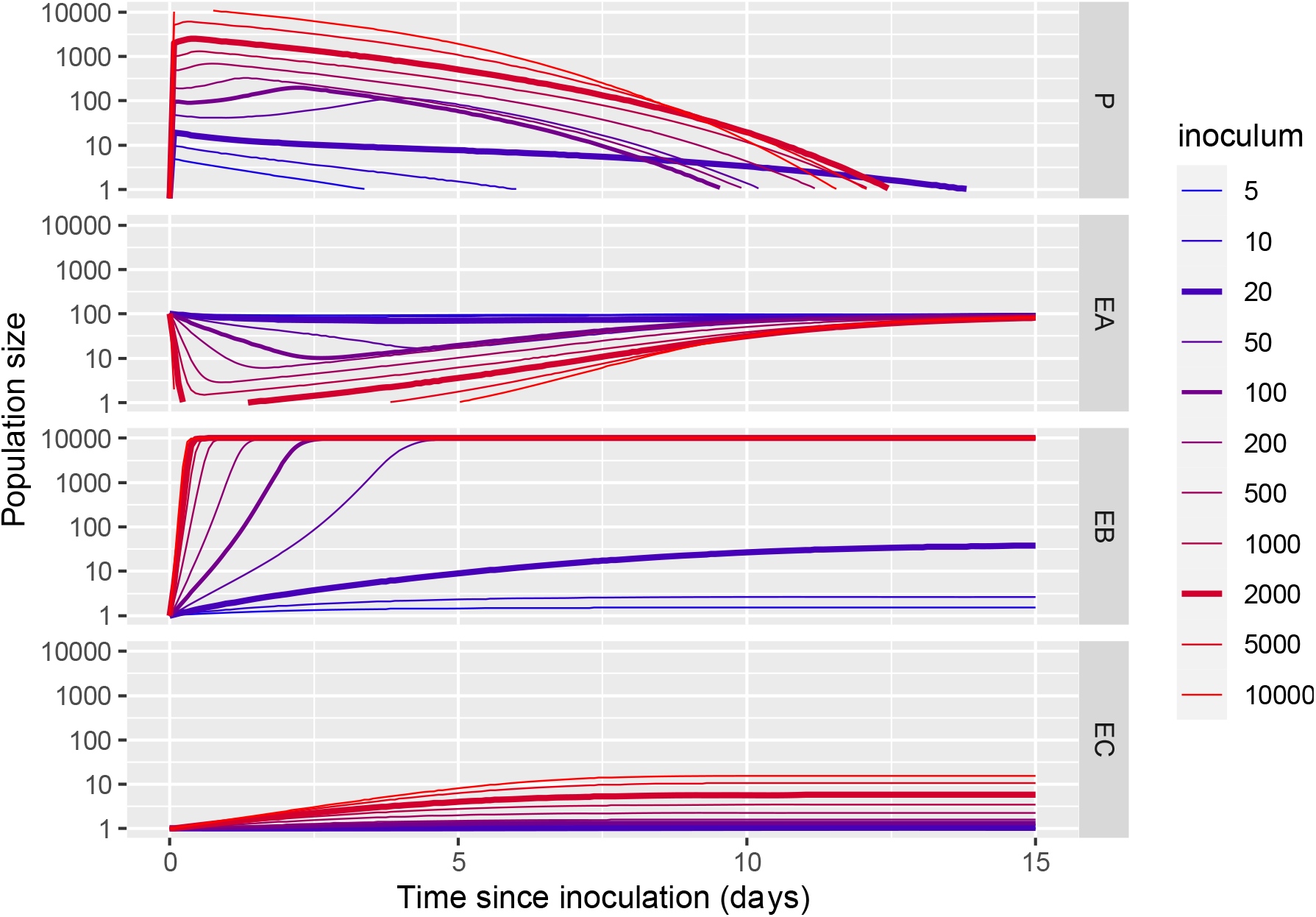
Time courses of parasite load (top row) and immune effectors of all three tiers (second, third and fourth rows) for varying inoculum sizes (doses). Intermediate inoculum sizes cause the longest infections due to how they interact with the series of defence tiers.

**Figure 1B:**
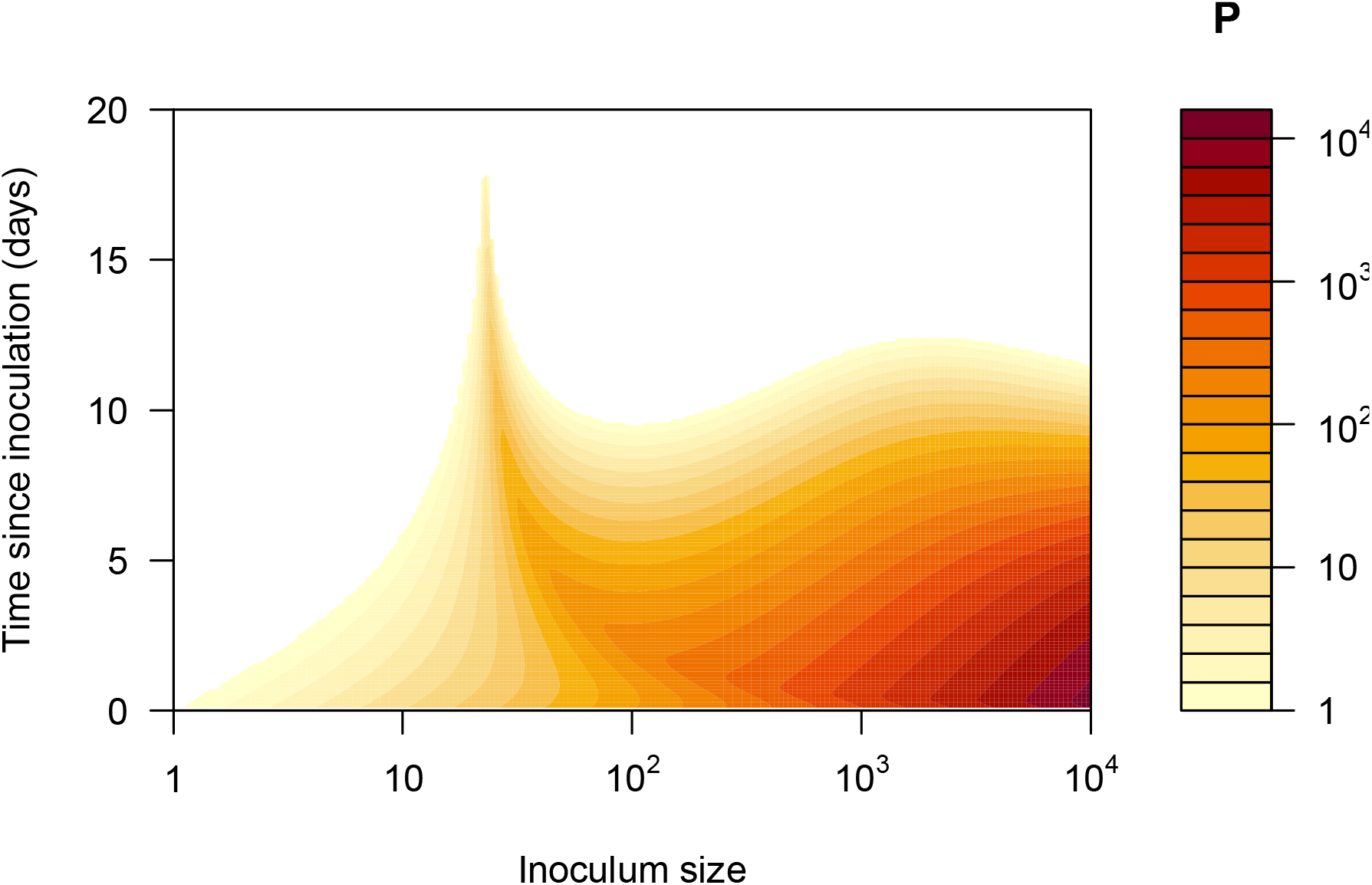
Time-course of parasite load for the varying doses displayed as a contour plot. (These are the same simulation data as shown in (A)). In this contour plot, it is more clearly visible that infection duration (in days since inoculation, on the y-axis) is longest at doses 20 and 2000, even if peak parasite load (colored according to heatmap P, at right) is highest in hosts receiving the largest dose. These simulations are based on the default parameterisation of our model (see Model Parameterisation and Scenarios and Supplementary Online Material.)

**Figure 1C:**
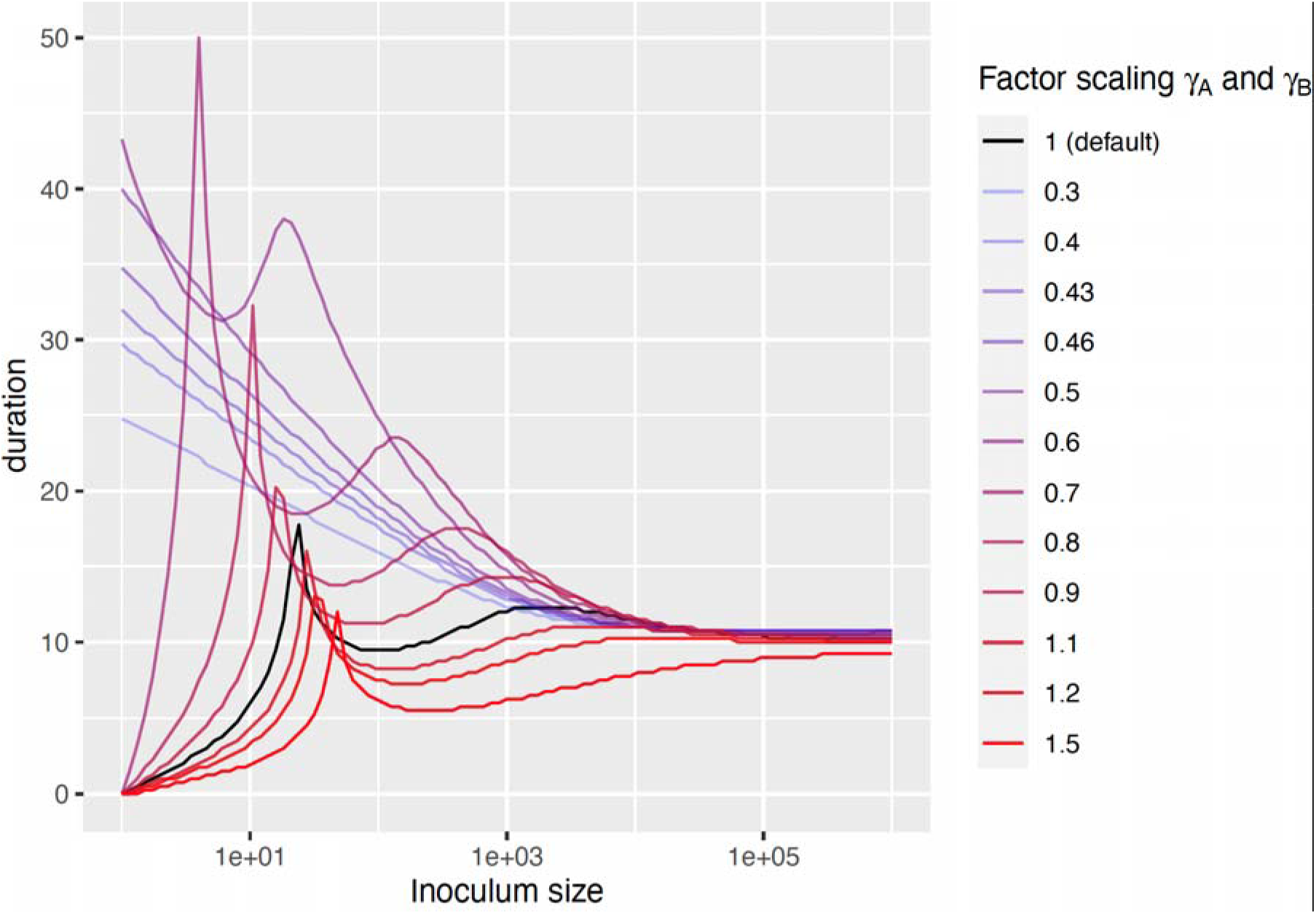
Infection duration versus inoculating dose for varying strengths of the first- and second-tier immune responses. The efficacy parameters of the first and second tier reponses, *γ*_*A*_/*γ*_*B*_, have been scaled to keep their relative strengths constant. The two peaks of infection duration are at lower doses for weaker immune responses, and for very weak responses they disappear completely, showing that the peaks in the relationship between duration and inoculum size are determined by the efficacy of the first- and second-tier defences.

The dynamics of parasites (**Figure 1A**) were non-monotonic: very low doses generated low peak parasite densities and short-lived infections; at higher doses, parasites persisted longer, with a peak in duration at the inoculum dose 20; at yet higher doses, infection duration decreased approximately two-fold, only to increase and peak again at an inoculum dose of 2000; beyond that, infection duration decreased again (**Figure 1B**). We also found that the multi-peaked relationship between inoculating dose and duration was robust to variation in the efficacy of barrier and induced defences, with more potent defences simply reducing the height of the peaks (**Figure 1C**).

We also tracked immune effector dynamics separately for the three tiers of defence. Dynamics of the height of the barrier (*E*_*A*_ in **Figure 1A**) exhibited non-monotonic dose-dependence, with low doses leaving the defence undepleted and high doses rapidly depleting the defence such that they rebounded quickly. Intermediate doses substantially depleted the barrier defences, which then rebounded more slowly than under high doses. By contrast, dynamics of the rapidly inducible (innate-like) and slowly-inducible (adaptive-like) defences varied in a more straight-forward fashion with dose. Innate immune effectors (*E*_*B*_ in **Figure 1A**) were induced increasingly rapidly with increasing dose, up to a maximum level of defence. Intermediate doses thereby induced middling innate defences. Adaptive immune effectors (*E*_*C*_ in **Figure 1A**) likewise exhibited straightforward dose dependence: both rates of induction and peaks of adaptive immune effectors increased monotonically with increasing parasite dose. Together, these results suggest that intermediate doses achieved long duration of infection by barely clearing the first or second tier defences and then inducing little further defence. In this way, such doses found a “wormhole” through 3-tiered host defence systems. (Hereby, we are referring to wormhole as described in the movie “Stargate”, rather than the one in old cupboards.)

We next systematically studied the peak parasite load and infection duration within 4 host types: “wild type” hosts with all 3 lines of defence intact; “barrier knockout” hosts lacking the first tier defence; “innate knockout” hosts lacking the rapidly inducible tier of defence; and “adaptive knockout” hosts lacking the slowly inducible tier of defence (**Figure 2**). To obtain sufficient resolution, we studied dose variation across six orders of magnitudes of inoculum size in each of these hosts.

**Figure 2:**
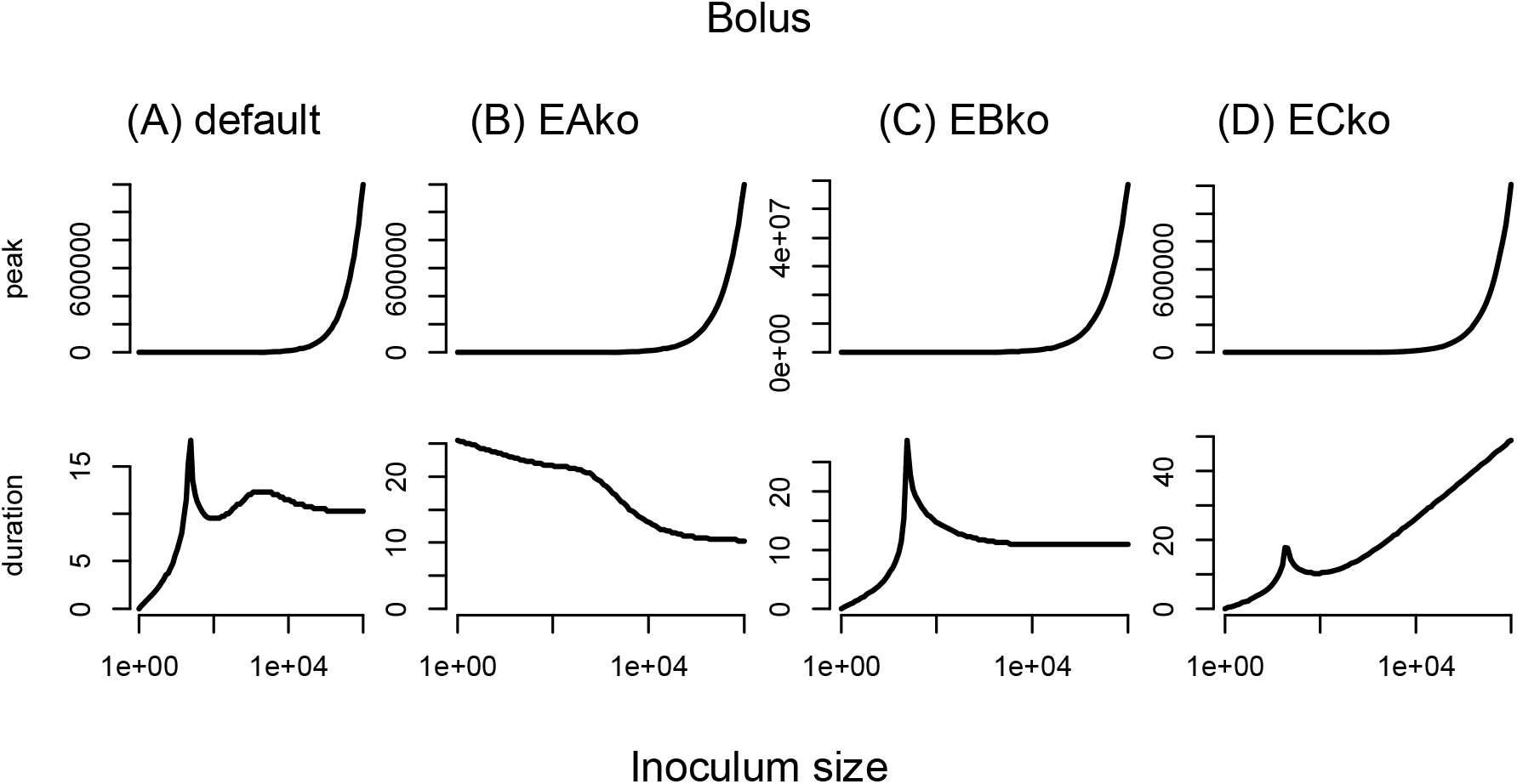
Peak parasite load (top row) and infection duration (bottom row) statistics for deterministic simulations across a range of inoculum sizes in 4 host types: (A) wildtype (“default”), (B) barrier knockouts (“EAko”), (C) second-tier (innate) knockouts (“EBko”), and (D) third-tier (adaptive) knockouts (“ECko”). Note the non-monotonic profiles of infection duration, especially in wildtype or innate-knockout hosts (“EBko”).

In wild-type “default” hosts (**Figure 2A**), we found that peak parasite load (top row) exhibited straightforward dose-dependence, increasing gradually with increasing dose. The <u>duration</u> of infection (second row), in contrast, exhibited non-monotonic dose dependence. Most interestingly, infection duration peaked twice across dose, as described above.

The three knockout host types provide insight into the causes of these patterns. Generally, for all types of knockouts (**Figures 2B-D**), the average duration of infections becomes longer across the range of doses compared to wild-type (see y-axis ranges).

Barrier knockout hosts exhibited simpler dose dependence than the wild-types, with peak parasite load (top row) increasing with increasing dose (**Figure 2B**). Most notably, however, infection duration (second row) did not display two peaks anymore but continuously decreased with dose. This decrease became more pronounced beyond a inoculation dose of approximately 500, echoing the second peak in the duration profile in the wild-type hosts (**Figure 2A**), while the first peak had disappeared. This change of pattern can be explained by the fact that knocking out barrier defences deletes the transition between first and second tier that is responsible for the first peak of the duration at low inoculum doses.

Knocking out innate responses (second tier), in contrast, makes the second peak in infection duration (second row) at high inoculum doses disappear (**Figure 2C**). The profiles of peak parasite load (top row) did not increase gradually anymore, but displayed jumps at an inoculum dose of 20, at which barrier responses are overcome and adaptive immune responses are triggered.

The patterns in adaptive immune knockouts are similar to those in innate immune knockouts: the second peak in the profile of infection duration versus dose (second row) disappears (**Figure 2D**). Unlike in innate immune knockouts, however, the infection duration does not continuously decrease with further dose escalation, but instead turns around and increases again. This difference in the patterns between innate versus adaptive immune knockouts may be due to the lower potency of innate compared to adaptive immune effectors.

These results show the link between the peaks in the profile of infection duration and tier transitions, and support a central role for tier transitions in explaining why intermediate doses of parasites lead to infections that persist longest.

Our final set of analyses of the deterministic system of equations aimed to assess the generality of the dose-dependent dynamics (**Figure 1**) and summary statistics (**Figure 2**) described thus far. We compared microparasites versus macroparasites (which, by definition, are non-replicating within a given host), and different patterns of exposure (a bit each day in a “trickle” versus all at once in a “bolus”); see **Supplementary Figures S1-S3**. For the macroparasite bolus exposures, the qualitative conclusion that duration depends non-monotonically on dose was the same as for microparasites, but details differed: e.g., intermediate doses generated a local, if not global, maximum for duration of infection even in barrier and innate response knockouts (**Figure S2**). Trickle exposures induced complexities, especially when parasites continued to arrive after considerable immune response induction and in hosts with a tier of defence knocked out (**Figures S1 and S3**). In no case was duration of infection monotonically associated with dose.

We next undertook stochastic simulations, to investigate how random variation in the processes hypothesized in our model would impact outcomes. We particularly focused on the response variable of infection duration, and we used finer-grain variation in inoculum size than before, but again across four orders of magnitude of dose. Consistent with the deterministic results, the stochastic results show non-monotonic dose dependence of infection duration with two peaks and tremendous variance associated with overcoming the first two tiers of defence. Again, doses intermediate between the extremes generated the most persistent infections (**Figure 3**).

**Figure 3:**
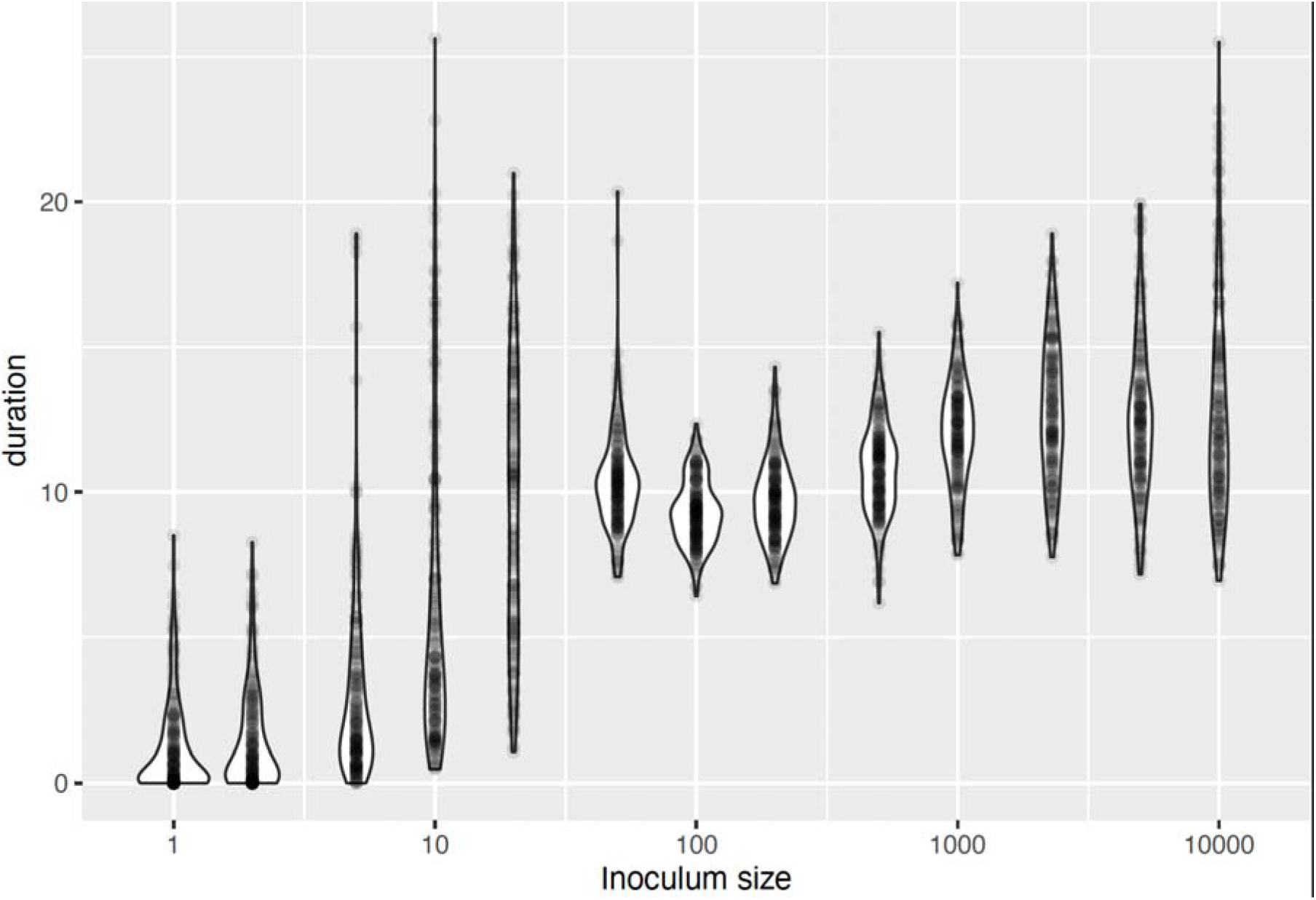
Infection duration versus inoculum size in stochastic simulations of the model, again showing non-monotonic changes in infection duration with increasing dose. Note that inoculum sizes of ∼50 that overcome the first-tier responses lead to less variable durations than inoculum sizes of 20.

Finally, we drew upon our stochastic formulation to investigate the probability of infection (infectivity), given inoculum size. We yet again observed non-monotonic, two-peaked dose-dependence, but time of observation emerged as crucial to the inference, as follows. Observed early after exposure, infection success appears to have a sigmoidal relationship to dose, with increasing success with dose up until a first peak. Observed later after exposure, hosts receiving high doses and thus more stimulation of inducible defence began to clear their infections, which generated a double-peaked relationship between dose and infection success (**Figure 4**). In other words, even infection success has two peaks at different intermediate doses, but detecting that relationship depends upon the timepoint at which hosts are observed and thus declared infected or not.

**Figure 4:**
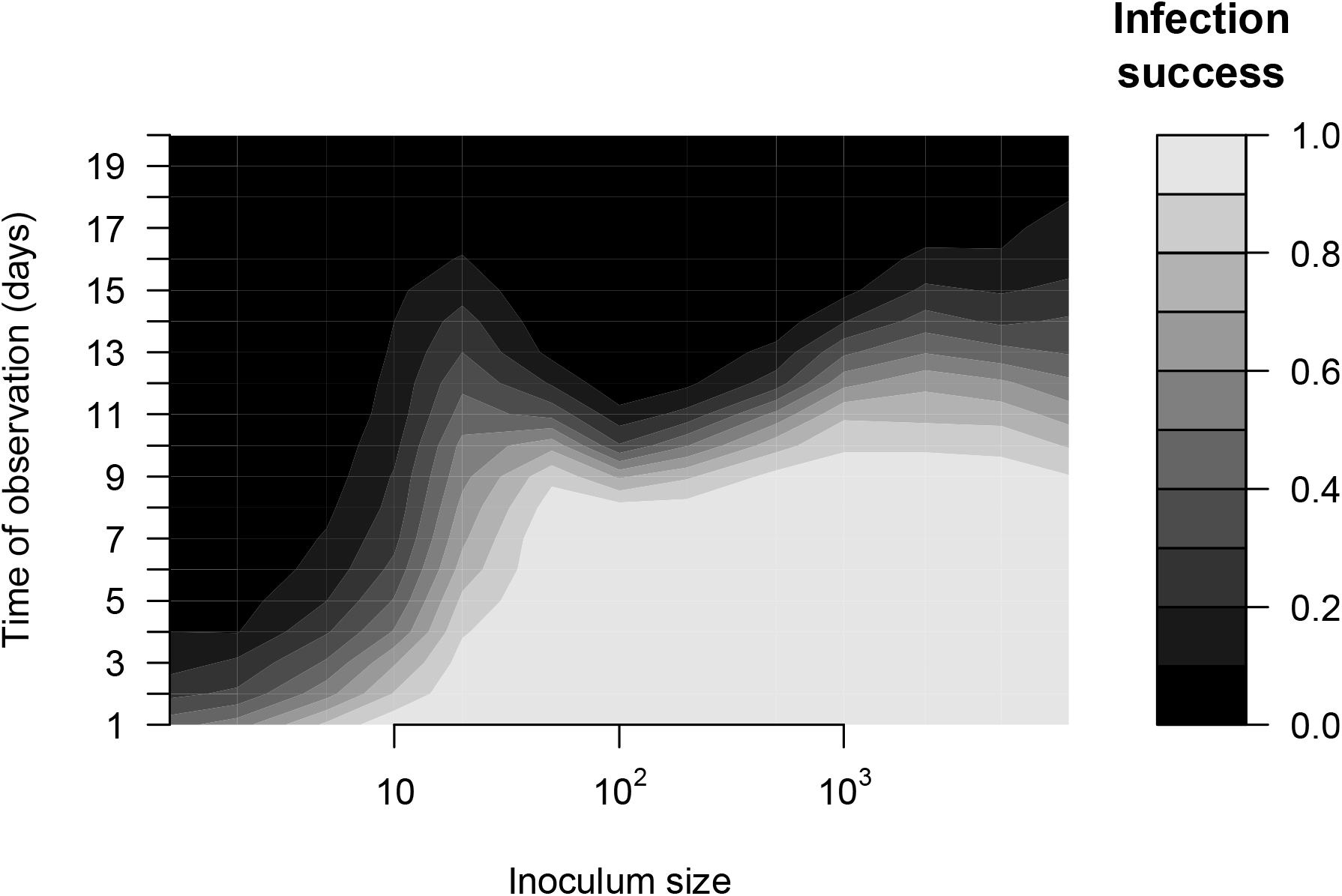
Infection success against inoculum size and time of observation. As expected, infection success increases overall with the inoculum size, and is higher when observed sooner. Note, however, the non-monotonic, double-peaked profile of infection success versus inoculum for observation times of ∼11 days.

Further simulations reveal that the stochastic version of our mathematical model can display bifurcating infection dynamics, such as those observed in (6, 28). **Figure S4** shows simulations of the time course of the parasite population in hosts, in which the second-tier innate responses are assumed to be knocked out. At the inoculum size that breaches the barrier response, the time courses are extremely variable leading to bi-modal infection duration distributions.

## Discussion

Immunoepidemiology links the dynamics of parasites within the host to their epidemiological spread. The two most essential parameters at this interface between immunology and epidemiology are exposure and susceptibility of hosts. Complicating immunoepidemiological analysis, however, is the fact that exposure and susceptibility are not independent of each other. Indeed, the number of parasites (or parasites per unit time (2)) to which hosts are <u>exposed</u> (here, inoculating dose) may in fact determine how <u>susceptible</u> the host is to infection *per se* (18-23), as well as susceptibility to chronic infection or severe symptoms (7, 8, 41-43). Studies of dose-dependence are therefore of applied relevance, and, in our view, are at the very heart of immunoepidemiology, because they also enable testing of basic science hypotheses such as the independent action of microbes (11, 12) and the optimal blend of constitutive and inducible defences (44). The value of studies of dose is greatly enhanced when physiological mechanisms are incorporated (24), as we have attempted here, with coarse but realistic incorporation of three tiers of host defence meeting varied inoculating doses.

Using a deterministic model that considers immune responses dynamically, we found that infection duration exhibited complex dose-dependence. Two intermediate inoculating doses generated infections of longest duration, while very low and very high doses, as well as doses between those chronicity-generating doses led to infections of shorter duration. We found that this is explained by certain doses being sufficient to clear the first or the second tier defences but then only slightly/slowly inducing subsequent tiers of effectors. This pattern is supported in at least some empirical systems (e.g., slow induction of antibodies by low doses of malaria parasites (45)).

Our findings thus suggest that parasites arriving at intermediate doses can thereby find “wormholes” to establishing and persisting in a host. Simulated knockout of barrier or innate defences turned the complex dose dependence with two peaks into simpler one, usually with a single peak of chronic infection associated with intermediate doses. Our simulations revealed threshold doses of approximately 20 and 2000 (for the parameters used) around which duration peaked in many cases. This corresponds to the dose necessary to clear either barrier or innate defences, and the variation suggests within-host feedbacks and stochasticity leading to high variation in trajectories of infection.

Before discussing such complexities, we note that our results, like others (2, 14, 26)}, are inconsistent with the hypothesis of independent action: i.e., we did not find the monotonic escalation of infection successwith increasing dose that is a hallmark of the multiplication of independent non-zero probabilities of success for each parasite (3-5). Our model thus suggests that the “one is enough” corollary of the independent action hypothesis, in which a single parasite has a considerable chance of establishing infection (46), is not general and depends upon the absence of diverse defences. Importantly, our model assumed no cooperativity to the parasites themselves. Instead, an “apparent cooperativity” arose due to dynamical interactions of the parasite population with each of the tiers of host defence.

Our central finding is that the first and second tiers of immune defences are crucial to both non-independent action of parasites and to chronicity-permitting “wormholes” in host defence. It is not new to suggest that constitutive defences are as important as inducible in explaining biotic interactions (47) including when defences against infection incur costs (e.g., (44, 48)). What is new here is the inference that this suite of defences may be what confers complex dose-dependence. This has been suggested by the importance of immune response growth terms that are independent of parasite load (25, 28, 29), but we extend this to barrier defence biology which may, in general, be central to predicting the probability and course of infection when host exposure varies.

Relevant barriers to infection are diverse (31), ranging from constitutive physical and chemical barriers (such as skin or stomach acid) through to continuously produced mucins, antimicrobial peptides (AMPs), or IgA on mucosal surfaces; such IgA can, for example, alter establishment of gut commensals (34) or prevent infection by pathogenic pneumococci (35). Dose-response studies of barrier defences are rare but have revealed, for example, that even barriers of intact skin and conjunctival mucosa are overcome at the highest doses of bacteria in genus *Leptospira* (49).

There are, of course, important caveats to acknowledge, but some of these also lead to insights. For example, we modeled just three coarse tiers, but in reality, within each coarse tier, there are finer grained tiers. Such granularity is evident in differential dependence of induced defences upon parasite load (e.g., (28, 29)) or immunisation rate (50) as well as varied innate defences of, for example, the lung (51, 52) or the diversity of T cell types that do not neatly fall in the classic innate vs adaptive categories (53). This array of finer tiers, including tiers of extremely rapidly induced defences, imply that “constitutive defences” actually are part of a continuum (31) of rates of defensive readiness. Our model revealed that three coarse tiers leaves vulnerabilities or “wormholes” around the barrier and innate defences; perhaps more realistic redundancies and finer-scaled tiers would mitigate such vulnerabilities. It is tempting to speculate that evolution has led to such redundancies and fine-scale tiers for the very reason that barriers open vulnerabilities. Nonetheless, plants (54) and animals such as cnidarians (55) are thought to have just two coarse tiers of defence (barrier & innate). It would be fascinating to investigate whether there are systematic differences in the relationship between inoculum dose and infection success or chronicity across hosts with two vs. three coarse tiers of defences. Our study suggests that in organisms with only two tiers of defence, there should be only one wormhole.

We look forward to further work in this area, both theoretical (to explore the general evolutionary defence logic) and empirical (to test the predictions). One testable prediction is the two-peaked dose-dependence in infection duration that should hold if there are three tiers of immune responses with two clearcut transitions among them. Dose-response studies with a wide range of doses could reveal such patterns.

In a second set of empirical follow-ups, one could also investigate the dose-dependences change in hosts with altered immune systems. Actual *in vivo* barrier knockouts analogous to our *in silico* barrier knockouts are rarely practical, but knockdowns may be: as in the work on leptospirosis of Gostic and colleagues (49), barrier defences like skin or mucosa could perhaps be experimentally eroded for dose-response studies across a broader array of infections. A further consideration is that empiricists designing dose-response studies are often logistically constrained to select just a few doses, and it may be challenging to identify a suitable range that will reveal any hidden nonlinearities, or even non-monotonicities. Indeed, the dose that is intermediate will vary hugely across parasite taxa, in ways that may map on to their life histories (56). The predictions generated here might thus best be tested via experimental approaches that track bottlenecks of founder bacteria in hosts deficient in immune defences of various speeds of activation and efficacy – e.g., barcoded but otherwise isogenic *Listeria monocytogenes* combined with host immune manipulation as in the elegant work of Zhang et al. (57).

Empirically-grounded theory has made recent progress in revealing rules of host-parasite interaction that accord with our findings here. These include stochasticity around inoculating dose thresholds or other early dynamics that lead to life or death for the host (6, 58). Another insight that has emerged from recent theory is that acute and chronic infections can emerge from the same set of equations so long as suitable feedback mechanisms (e.g., between parasite growth and induced immune defences, or among different induced immune responses are built in (59, 60)). Our work here is in that vein but adds that complex dose-dependence of the probability and duration of infection may arise from the interplay of host cells and parasite populations. Insights from within-host dynamics arguably have major implications for both immunoepidemiology of diverse infections and the evolution of defence systems.

## Acknowledgements

ALG thanks ETH Zürich for sabbatical fellowship support, and both authors thank Aaron King, Dimitri Diavatopoulos, and Paul Schmid-Hempel for thought-provoking discussions.

## Supplementary figures

**Figure S1:**
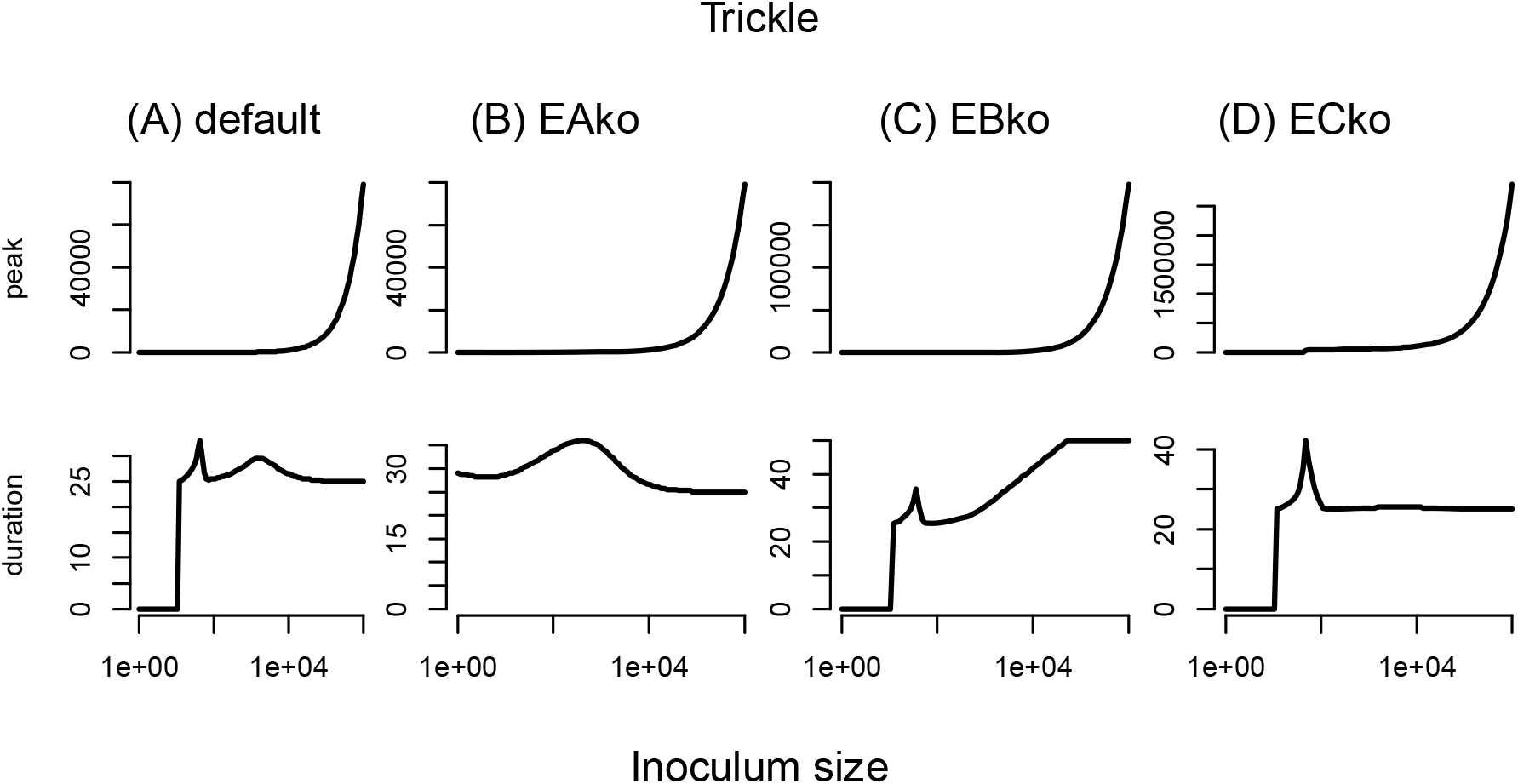
Peak parasite load (top row) and infection duration (bottom row) statistics for deterministic simulations across a range of inoculum sizes in 4 host types: (A) wildtype (“default”), (B) barrier knockouts (“EAko”), (C) second-tier knockouts (“EBko”), and (D) third-tier knockouts (“ECko”). Here, all doses arrive slowly over time, in a trickle rather than bolus inoculation.

**Figure S2:**
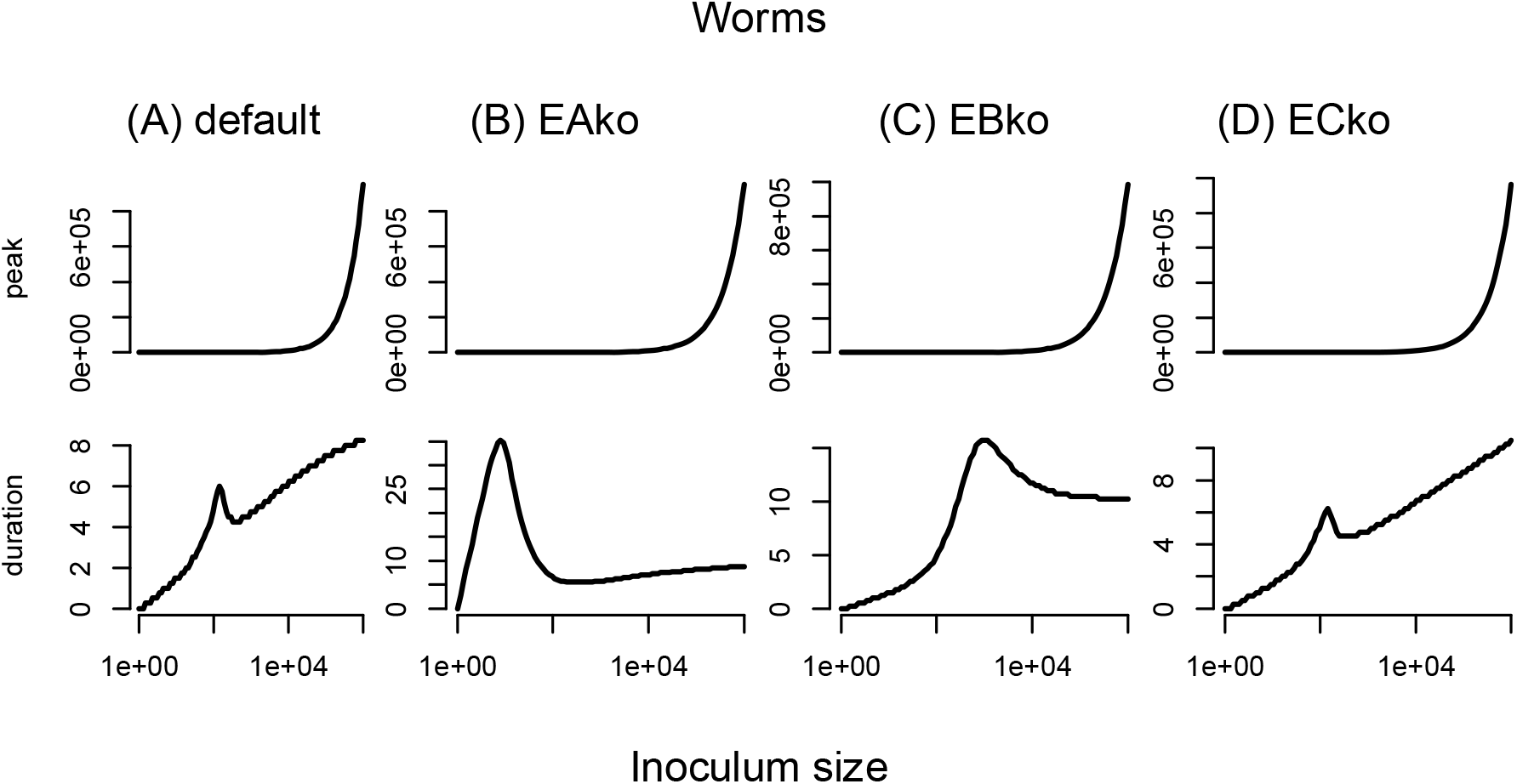
Peak parasite load (top row) and infection duration (bottom row) statistics for deterministic simulations across a range of inoculum sizes in 4 host types: (A) wildtype (“default”), (B) barrier knockouts (“EAko”), (C) second-tier knockouts (“EBko”), and (D) third-tier knockouts (“ECko”). Here, all parasites are non-replicating macroparasites, shorthanded here as worms, that arrive all at once in a bolus inoculation.

**Figure S3:**
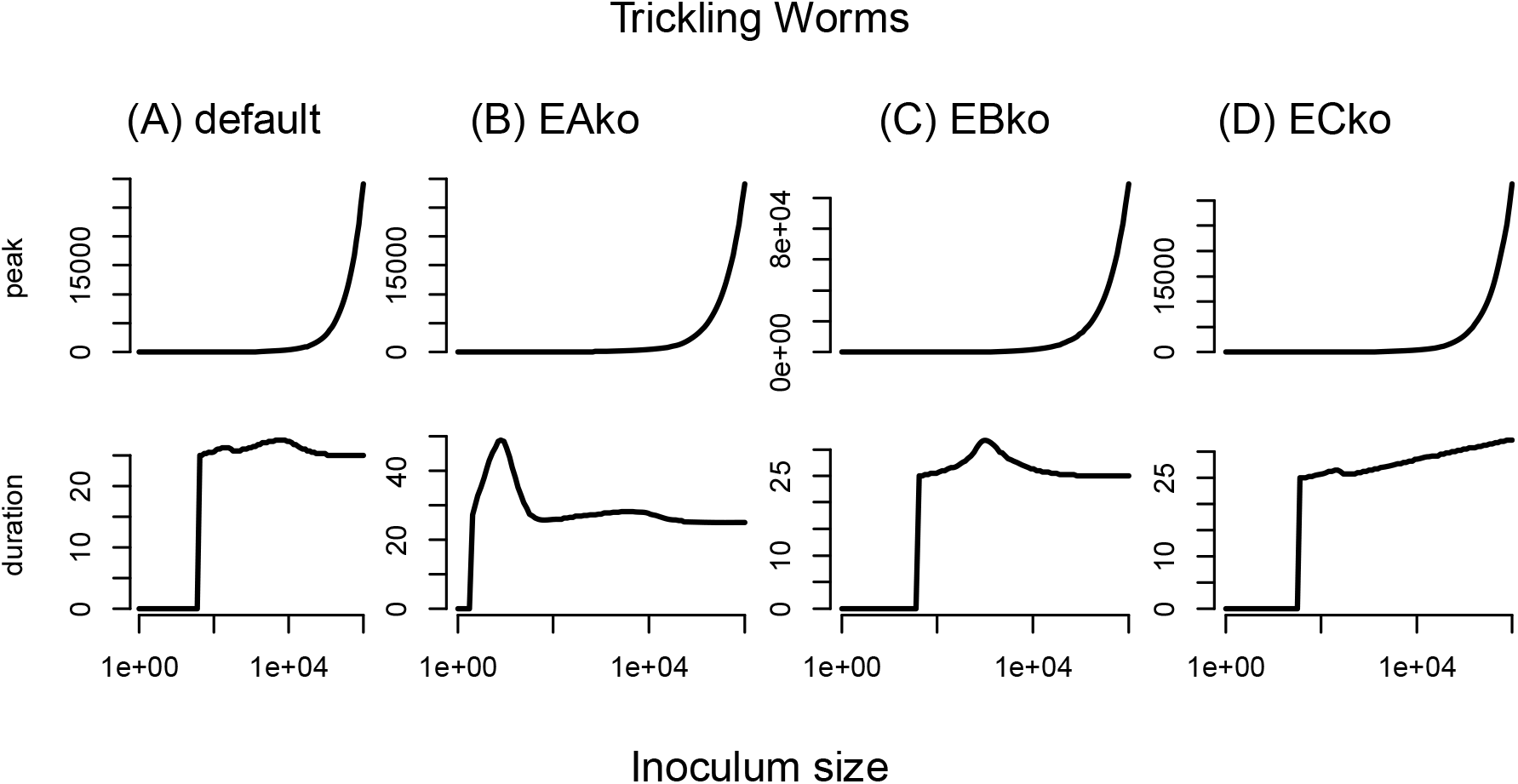
Peak parasite load (top row) and infection duration (bottom row) statistics for deterministic simulations across a range of inoculum sizes in 4 host types: (A) wildtype (“default”), (B) barrier knockouts (“EAko”), (C) second-tier knockouts (“EBko”), and (D) third-tier knockouts (“ECko”). Here, all parasites are non-replicating macroparasites, shorthanded here as worms, that arrive slowly in a trickle inoculation.

**Figure S4:**
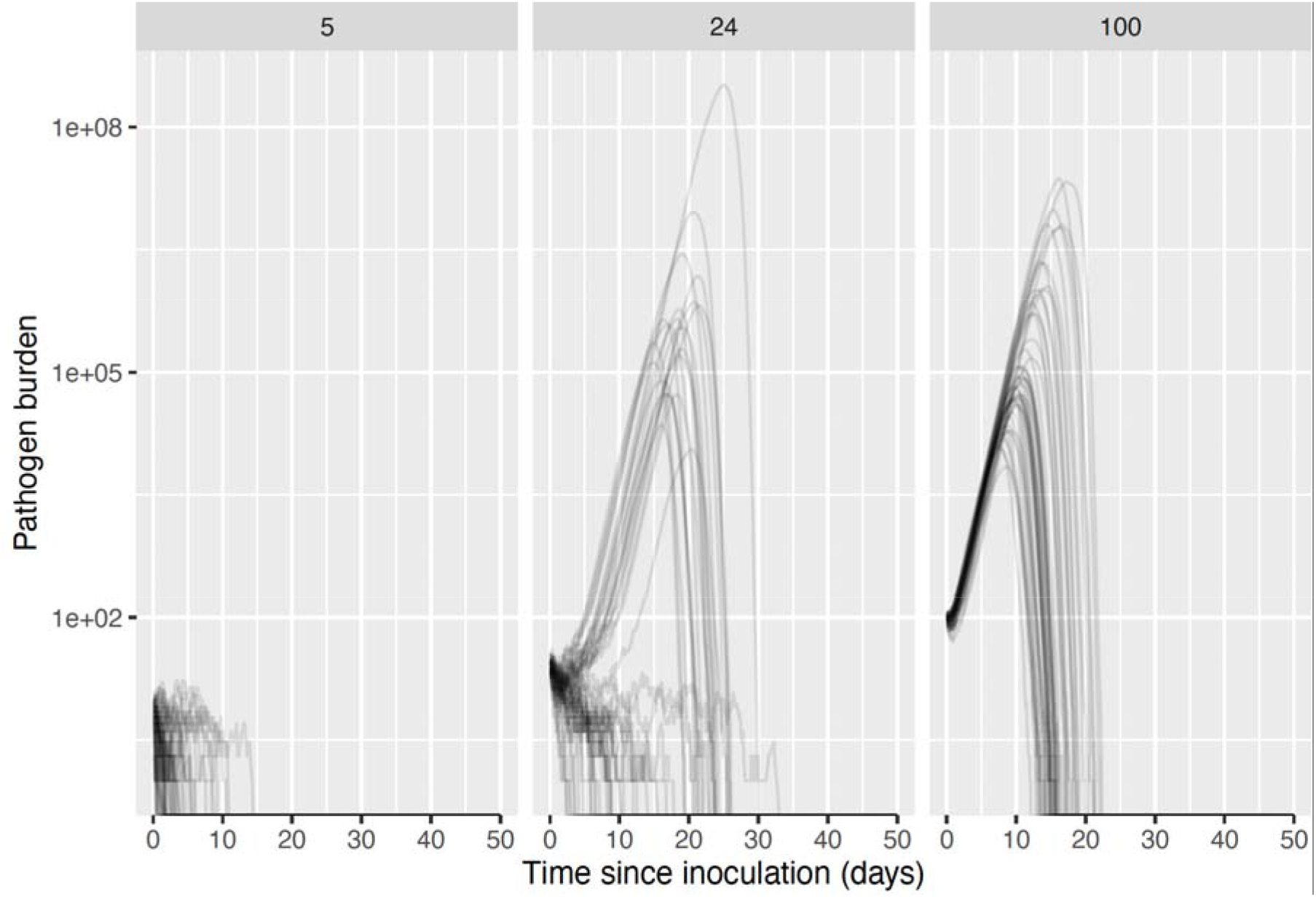
Stochastic simulations of the time course of the parasite population in hosts in which the second-tier innate responses are assumed to be knocked out.

## References

1. Frank SA. Immune response to parasitic attack: evolution of a pulsed character. J Theor Biol. 2002;219(3):281–90.

2. Pujol JM, Eisenberg JE, Haas CN, Koopman JS. The effect of ongoing exposure dynamics in dose response relationships. PLoS Comput Biol. 2009;5(6):e1000399.

3. Zwart MP, Hemerik L, Cory JS, de Visser JA, Bianchi FJ, Van Oers MM, et al. An experimental test of the independent action hypothesis in virus-insect pathosystems. Proc Biol Sci. 2009;276(1665):2233–42.

4. Zwart MP, Elena SF. Testing the independent action hypothesis of plant pathogen mode of action: a simple and powerful new approach. Phytopathology. 2015;105(1):18–25.

5. McKenney DG, Kurath G, Wargo AR. Characterization of infectious dose and lethal dose of two strains of infectious hematopoietic necrosis virus (IHNV). Virus Res. 2016;214:80–9.

6. Duneau D, Ferdy JB, Revah J, Kondolf H, Ortiz GA, Lazzaro BP, et al. Stochastic variation in the initial phase of bacterial infection predicts the probability of survival in D. melanogaster. Elife. 2017;6.

7. Ebert D, Zschokke-Rohringer CD, Carius HJ. Dose effects and density-dependent regulation of two microparasites of Daphnia magna. Oecologia. 2000;122(2):200–9.

8. Paterson S, Wilkes C, Bleay C, Viney ME. Immunological responses elicited by different infection regimes with Strongyloides ratti. PLoS One. 2008;3(6):e2509.

9. Wunder EA, Jr., Figueira CP, Santos GR, Lourdault K, Matthias MA, Vinetz JM, et al. Real-Time PCR Reveals Rapid Dissemination of Leptospira interrogans after Intraperitoneal and Conjunctival Inoculation of Hamsters. Infect Immun. 2016;84(7):2105–15.

10. Hamilton KA, Weir MH, Haas CN. Dose response models and a quantitative microbial risk assessment framework for the Mycobacterium avium complex that account for recent developments in molecular biology, taxonomy, and epidemiology. Water Res. 2017;109:310–26.

11. Halvorson HO. The effect of chance on the mortality of experimentally infected animals. J Bacteriol. 1935;30:330.

12. Druett HA. Bacterial invasion. Nature. 1952;170(4320):288.

13. Moxon ER, Murphy PA. Haemophilus influenzae bacteremia and meningitis resulting from survival of a single organism. Proc Natl Acad Sci U S A. 1978;75(3):1534–6.

14. Shao X, Levin B, Nemenman I. Single variant bottleneck in the early dynamics of H. influenzae bacteremia in neonatal rats questions the theory of independent action. Phys Biol. 2017;14(4):045004.

15. Cornforth DM, Matthews A, Brown SP, Raymond B. Bacterial Cooperation Causes Systematic Errors in Pathogen Risk Assessment due to the Failure of the Independent Action Hypothesis. PLoS Pathog. 2015;11(4):e1004775.

16. Cornforth DM, Popat R, McNally L, Gurney J, Scott-Phillips TC, Ivens A, et al. Combinatorial quorum sensing allows bacteria to resolve their social and physical environment. Proc Natl Acad Sci U S A. 2014;111(11):4280–4.

17. Mony BM, MacGregor P, Ivens A, Rojas F, Cowton A, Young J, et al. Genome-wide dissection of the quorum sensing signalling pathway in Trypanosoma brucei. Nature. 2014;505(7485):681–5.

18. Ben-Ami F, Ebert D, Regoes RR. Pathogen dose infectivity curves as a method to analyze the distribution of host susceptibility: a quantitative assessment of maternal effects after food stress and pathogen exposure. Am Nat. 2010;175(1):106–15.

19. Ben-Ami F, Regoes RR, Ebert D. A quantitative test of the relationship between parasite dose and infection probability across different host-parasite combinations. Proc Biol Sci. 2008;275(1636):853–9.

20. Regoes RR, Hottinger JW, Sygnarski L, Ebert D. The infection rate of Daphnia magna by Pasteuria ramosa conforms with the mass-action principle. Epidemiol Infect. 2003;131(2):957–66.

21. Regoes RR. The role of exposure history on HIV acquisition: insights from repeated low-dose challenge studies. PLoS Comput Biol. 2012;8(11):e1002767.

22. Langwig KE, Wargo AR, Jones DR, Viss JR, Rutan BJ, Egan NA, et al. Vaccine Effects on Heterogeneity in Susceptibility and Implications for Population Health Management. mBio. 2017;8(6).

23. Dwyer G, Elkinton JS, Buonaccorsi JP. Host heterogeneity in susceptibility and disease dynamics: tests of a mathematical model. Am Nat. 1997;150(6):685–707.

24. Haas CN. Microbial dose response modeling: past, present, and future. Environ Sci Technol. 2015;49(3):1245–59.

25. Althouse BM, Hanley KA. The tortoise or the hare? Impacts of within-host dynamics on transmission success of arthropod-borne viruses. Philos Trans R Soc Lond B Biol Sci. 2015;370(1675).

26. Li Y, Handel A. Modeling inoculum dose dependent patterns of acute virus infections. J Theor Biol. 2014;347:63–73.

27. Pilyugin SS, Antia R. Modeling immune responses with handling time. Bull Math Biol. 2000;62(5):869–90.

28. Tate AT, Graham AL. Dissecting the contributions of time and microbe density to variation in immune gene expression. Proc Biol Sci. 2017;284(1859).

29. Jent D, Perry A, Critchlow J, Tate AT. Natural variation in the contribution of microbial density to inducible immune dynamics. Mol Ecol. 2019;28(24):5360–72.

30. Levin BR, Antia R. Why we don’t get sick: the within-host population dynamics of bacterial infections. Science. 2001;292(5519):1112–5.

31. Paludan SR, Pradeu T, Masters SL, Mogensen TH. Constitutive immune mechanisms: mediators of host defence and immune regulation. Nat Rev Immunol. 2021;21(3):137–50.

32. Fenton A, Perkins SE. Applying predator-prey theory to modelling immune-mediated, within-host interspecific parasite interactions. Parasitology. 2010;137(6):1027–38.

33. Metcalf CJ, Graham AL, Huijben S, Barclay VC, Long GH, Grenfell BT, et al. Partitioning regulatory mechanisms of within-host malaria dynamics using the effective propagation number. Science. 2011;333(6045):984–8.

34. Rollenske T, Burkhalter S, Muerner L, von Gunten S, Lukasiewicz J, Wardemann H, et al. Parallelism of intestinal secretory IgA shapes functional microbial fitness. Nature. 2021.

35. Binsker U, Lees JA, Hammond AJ, Weiser JN. Immune exclusion by naturally acquired secretory IgA against pneumococcal pilus-1. J Clin Invest. 2020;130(2):927–41.

36. Mak TW, Penninger JM, Ohashi PS. Knockout mice: a paradigm shift in modern immunology. Nat Rev Immunol. 2001;1(1):11–9.

37. Team RC. R: A language and environment for statistical computing Vienna, Austria: R Foundation for Statistical Computing; 2021 [Available from: https://www.R-project.org/.

38. Soetaert K, Petzoldt T, Setzer RW. Solving differential equations in r: Package desolve. Journal of Statistical Software. 2010;33(9):1–25.

39. Johnson P. Tau-leaping stochastic simulation: package ‘adaptivetau’: R Foundation for Statistical Computing; 2019 [Available from: https://CRAN.R-project.org/package=adaptivetau.

40. Cao Y, Gillespie DT, Petzold LR. Adaptive explicit-implicit tau-leaping method with automatic tau selection. Journal of Chemical Physics. 2007;126:224101.

41. Scott ME. Heligmosomoides polygyrus (Nematoda): susceptible and resistant strains of mice are indistinguishable following natural infection. Parasitology. 1991;103 Pt 3:429–38.

42. Scott ME. High transmission rates restore expression of genetically determined susceptibility of mice to nematode infections. Parasitology. 2006;132(Pt 5):669–79.

43. Chan MS, Medley GF, Jamison D, Bundy DA. The evaluation of potential global morbidity attributable to intestinal nematode infections. Parasitology. 1994;109 (Pt 3):373–87.

44. Hamilton R, Siva-Jothy M, Boots M. Two arms are better than one: parasite variation leads to combined inducible and constitutive innate immune responses. Proc Biol Sci. 2008;275(1637):937–45.

45. Fairlie-Clarke KJ, Hansen C, Allen JE, Graham AL. Increased exposure to Plasmodium chabaudi antigens sustains cross-reactivity and avidity of antibodies binding Nippostrongylus brasiliensis: dissecting cross-phylum cross-reactivity in a rodent model. Parasitology. 2015;142(14):1703–14.

46. Zwart MP, Daros JA, Elena SF. One is enough: in vivo effective population size is dose-dependent for a plant RNA virus. PLoS Pathog. 2011;7(7):e1002122.

47. Tollrian R, Harvell CD, editors. The Ecology and Evolution of Inducible Defences. Princeton, New Jersey: Princeton University Press; 1999.

48. Cressler CE, Graham AL, Day T. Evolution of hosts paying manifold costs of defence. Proc Biol Sci. 2015;282(1804):20150065.

49. Gostic KM, Wunder EA, Jr., Bisht V, Hamond C, Julian TR, Ko AI, et al. Mechanistic dose-response modelling of animal challenge data shows that intact skin is a crucial barrier to leptospiral infection. Philos Trans R Soc Lond B Biol Sci. 2019;374(1782):20190367.

50. Cirelli KM, Carnathan DG, Nogal B, Martin JT, Rodriguez OL, Upadhyay AA, et al. Slow Delivery Immunization Enhances HIV Neutralizing Antibody and Germinal Center Responses via Modulation of Immunodominance. Cell. 2019;177(5):1153–71 e28.

51. Iwasaki A, Foxman EF, Molony RD. Early local immune defences in the respiratory tract. Nat Rev Immunol. 2017;17(1):7–20.

52. Whitsett JA, Alenghat T. Respiratory epithelial cells orchestrate pulmonary innate immunity. Nat Immunol. 2015;16(1):27–35.

53. Mayassi T, Barreiro LB, Rossjohn J, Jabri B. A multilayered immune system through the lens of unconventional T cells. Nature. 2021;595(7868):501–10.

54. Yuan M, Jiang Z, Bi G, Nomura K, Liu M, Wang Y, et al. Pattern-recognition receptors are required for NLR-mediated plant immunity. Nature. 2021;592(7852):105–9.

55. Parisi MG, Parrinello D, Stabili L, Cammarata M. Cnidarian Immunity and the Repertoire of Defence Mechanisms in Anthozoans. Biology (Basel). 2020;9(9).

56. Schmid-Hempel P, Frank SA. Pathogenesis, virulence, and infective dose. PLoS Pathog. 2007;3(10):1372–3.

57. Zhang T, Abel S, Abel Zur Wiesch P, Sasabe J, Davis BM, Higgins DE, et al. Deciphering the landscape of host barriers to Listeria monocytogenes infection. Proc Natl Acad Sci U S A. 2017;114(24):6334–9.

58. Tate AT, Andolfatto P, Demuth JP, Graham AL. The within-host dynamics of infection in trans-generationally primed flour beetles. Mol Ecol. 2017;26(14):3794–807.

59. van Leeuwen A, Budischak SA, Graham AL, Cressler CE. Parasite resource manipulation drives bimodal variation in infection duration. Proc Biol Sci. 2019;286(1902):20190456.

60. Ellner SP, Buchon N, Dorr T, Lazzaro BP. Host-pathogen immune feedbacks can explain widely divergent outcomes from similar infections. Proc Biol Sci. 2021;288(1951):20210786.

